# High-content microscopy and machine learning characterize a cell morphology signature of *NF1* genotype in Schwann cells

**DOI:** 10.1101/2024.09.11.612546

**Authors:** Jenna Tomkinson, Cameron Mattson, Michelle Mattson-Hoss, Herb Sarnoff, Stephanie J. Bouley, James A. Walker, Gregory P. Way

## Abstract

Neurofibromatosis type 1 (NF1) is a multi-system, autosomal dominant genetic disorder driven by the systemic loss of the NF1 protein neurofibromin. Loss of neurofibromin in Schwann cells is particularly detrimental, as the acquisition of a ‘second-hit’ (e.g., complete loss of *NF1*) can lead to the development of plexiform neurofibromas (pNF). pNFs are painful, disfiguring tumors with an approximately 1 in 5 chance of sarcoma transition. Selumetinib and mirdametinib are currently the only medicines approved by the U.S. Food and Drug Administration (FDA) for the treatment of pNFs. This motivates the need to develop new therapies, either derived to treat *NF1* haploinsufficiency or complete loss of *NF1* function. To identify new therapies, we need to understand the impact neurofibromin has on Schwann cells. Here, we aimed to characterize differences in high-content microscopy in neurofibromin-deficient Schwann cells. We applied a fluorescence microscopy assay (called Cell Painting) to an isogenic pair of Schwann cell lines (derived from ipn02.3 2λ), one of wildtype genotype (*NF1*^*+/+*^) and one of *NF1* null genotype (*NF1*^*-/-*^). We modified the canonical Cell Painting assay to mark four organelles/subcellular compartments: nuclei, endoplasmic reticulum, mitochondria, and F-actin. We utilized CellProfiler to perform quality control, illumination correction, segmentation, and cell morphology feature extraction. We segmented 20,680 *NF1* wildtype and null cells, measured 894 significant cell morphology features representing various organelle shapes and intensity patterns, and trained a logistic regression machine learning model to predict the *NF1* genotype of single Schwann cells. The machine learning model had high performance, with training and testing data yielding a balanced accuracy of 0.85 and 0.80, respectively. However, when applied to a new pair of Schwann cells, the model’s balanced accuracy dropped to 0.5, which is no better than random chance. This performance decline appears to result from morphology differences introduced by non-biological factors (cloning procedures, origin of parental cell line, and CRISPR procedures) of the second cell line pair. We plan to improve upon this preliminary model by refining the *NF1* morphology signature using a broader panel of Schwann cell lines. Our goal is to apply this enhanced signature in large-scale drug screens of *NF1*-deficient cells to identify candidate therapeutic agents that specifically reverse the disease-associated morphology. Ultimately, we aim to identify agents that restore NF1 patient-derived Schwann cells to a phenotype resembling the *NF1* wild-type and healthier state.

## Introduction

Neurofibromatosis type 1 (NF1) is a haploinsufficient, autosomal dominant genetic condition that impacts approximately 1 in 3,000 individuals worldwide.^1–3^ NF1 results from mutations in the *NF1* gene, with approximately half the affected individuals developing the disorder due to *de novo NF1* pathogenic variants.^4^ The *NF1* gene encodes the protein neurofibromin, a GTPase-activating protein that negatively regulates Ras. Loss of NF1 results in increased Ras signaling effector and downstream pathways such as MAPK and mTOR signaling, which can cause unchecked cell proliferation.^5^ Benign and malignant tumor development is connected to a “second hit” somatic mutation in the *NF1* gene, which results from a loss of heterozygosity (LOH) and additional mutations.^6,7^ Generally, tumors are initiated by neurofibromin-deficient (*NF1*^*-/-*^) Schwann cells, and driven by *NF1* haploinsufficient microenvironments.^1,2^

Over 99% of NF1 patients develop cutaneous neurofibromas (cNFs), benign skin tumors that significantly decrease quality of life.^8,9^ Furthermore, about 30-50% of NF1 patients develop plexiform neurofibromas (pNFs), benign tumors that grow from nerves and are often painful and disfiguring. When surgically resected, 50% of pNFs regrow within 10 years.^10^ Another 8-15% of NF1 patients have pNFs that transform into malignant peripheral nerve sheath tumors (MPNSTs), which are rare sarcomas that have a five-year survival rate that ranges greatly from patient to patient but can be as low as 16% or as high as 62%.^11,12^ There are currently only two FDA-approved treatments for NF1 patients, selumetinib and mirdametinib. Both are MEK1/2 inhibitors that reduce pNF growth by suppressing MAPK signaling to reduce cell proliferation.^13,14^ However, both treatments are only approved to treat pNFs (not cNFs nor MPNSTs), and recent evidence shows selumetinib has limited efficacy with only 30-50% shrinkage in about half of patients, and many patients experience significant adverse events.^15,16^

We previously proposed that therapeutically elevating neurofibromin activity in *NF1* haploinsufficient Schwann cell environments will slow or reverse tumor development and other NF1 manifestations.^17^ One way to identify treatments that therapeutically elevate neurofibromin is to screen many drugs that cause the desired effect. We and others have performed NF1 drug screening applications previously, but they either use cell types unrelated to NF1 and/or measure only simple readouts like cell viability or other endpoints that will not prioritize a drug that reverses specific phenotypes.^18–20^ A revitalized approach, called phenotypic drug discovery, applies modern computational and microscopy tools to prioritize drugs based on their phenotypic effects on relevant cell models.^21^ However, in order to pursue phenotypic drug discovery, we must first confirm feasibility and select an appropriate assay to identify a sensitive and specific phenotypic signature. Therefore, we aimed to demonstrate proof of concept that the under-utilized approach known as high-content microscopy can distinguish *NF1* genotype in Schwann cells. High-content microscopy, which enables scientists to extract rich phenotypic information directly from microscopy images of cells, has greatly evolved over the past twenty years, and it is now poised to accelerate its impact on many different diseases, including NF1.^22^ Many studies have quantified molecular differences in Schwann cells with and without NF1^23–25^; however, few studies have quantified phenotypic differences. Kim et al., 1995, for example, analyzed phase-contrast micrographs of Schwann cells from each genotype, identifying that complete neurofibromin loss (*NF1*^*-/-*^) resulted in elongated cell processes and bright-refractile cell bodies as compared to wildtype cells (*NF1*^*+/+*^).^26^ While this study identified qualitative morphological differences, it was conducted through manual observation instead of an unbiased automatic system of quantitative data analysis.

Here, we utilized modern advances in high-content microscopy to quantify cellular phenotypic variations at the single-cell level. We employed a modified version of the multiplex fluorescence microscopy assay called Cell Painting to mark four different organelles/subcellular compartments (nuclei, endoplasmic reticulum (ER), mitochondria, and F-actin cytoskeleton).^27^ We used CellProfiler^28^ to segment and measure quantitative cell morphology features in single cells.^29^ We applied advanced image analysis and a machine learning pipeline to identify significant single-cell morphology differences in isogenic Schwann cells (derived from an ipn02.3 2λ parental line) with different *NF1* genotypes (*NF1*^*+/+*^ *and NF1*^*-/-*^). Our high-content microscopy and machine learning approach predicts *NF1* genotypes with high performance, providing proof of concept that we can use phenotypic drug discovery to identify treatments that reverse *NF1*-deficient phenotypes. Moreover, we identified morphology features of the endoplasmic reticulum (ER) and F-actin cytoskeleton that strongly differentiate *NF1* genotypes. However, when applying our pretrained model to a different isogenic pair of Schwann cells derived from the same parental line but with an alternative approach, we observed performance no greater than random chance. This highlights both the model’s sensitivity to subtle signatures and the importance of precise experimental procedures in isolating a high-content microscopy signature of *NF1* genotype in Schwann cells. This work sets the stage for future studies where we will refine the *NF1* morphology signature using a broader panel of Schwann cell lines (including primary Schwann cells), and apply high-throughput phenotypic drug discovery to identify therapeutic agents that elevate wildtype neurofibromin levels, potentially slowing or reversing tumor development in NF1 patients.

## Results

### Processing single-cell morphological profiles of Schwann cells

We applied our modified Cell Painting assay to mark nuclei, ER, mitochondria, and F-actin of hTERT ipn02.3 2λ Schwann cells (*NF1*^*+/+*^) and a CRISPR engineered isogenic null cell (*NF1*^*-/-*^) (**Figure 1A**).^30^ We call these lines A3 and C04, respectively, as previously described.^31^ We confirmed loss of neurofibromin in C04 via Western blot (**Supplementary Figure 1**). We employed image analysis and machine learning pipelines to extract, process, and analyze high-dimensional single-cell morphology features to identify an *NF1* genotype signature (**Figure 1B**). For our image analysis pipeline, we used CellProfiler to perform illumination correction to retrospectively correct uneven lighting across the images, including vignetting.^32^ We then segmented nuclei, whole cells, and cytoplasm and extracted a total of 2,313 morphology features. We then applied image-based profiling^33^, a method that formats morphological profiles to evaluate differences among cell populations. Specifically, we applied CytoTable to curate single-cell profiles^34^ and pycytominer to annotate, normalize, select features, and aggregate these data to analyze downstream with machine learning.^35^ We also used coSMicQC^36^ to perform single-cell quality control (QC), which filtered 9.2% of cells across all plates (**Supplementary Figure 2**). To isolate morphology differences in non-dividing cells (we expect increased cell proliferation in *NF1* Null Schwann cells^26^), our filtering included removing cells likely undergoing mitosis.^37^

**Figure 1.**
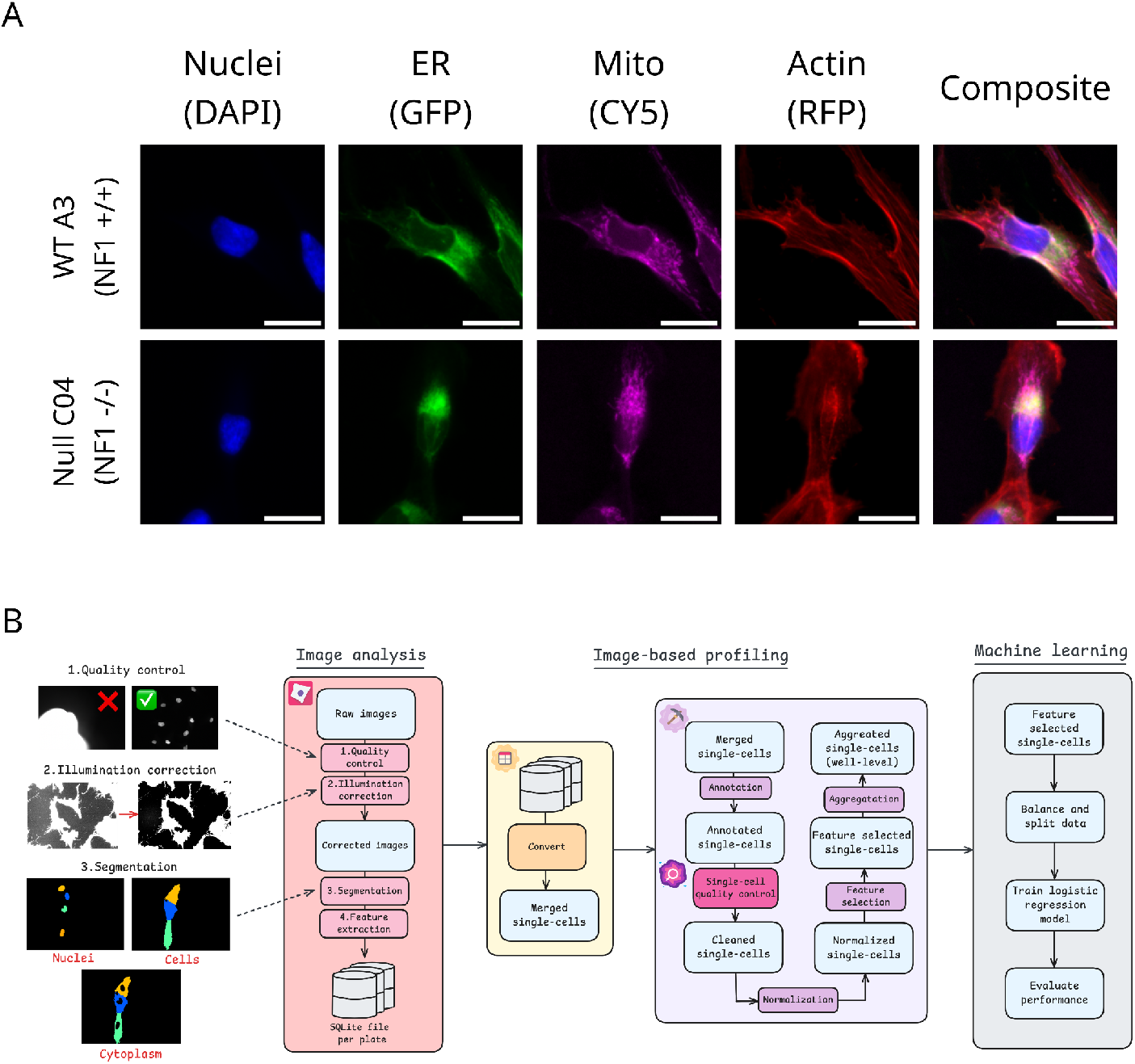
Overview of our Cell Painting and data analysis workflow. (**A**) Example image montages of the Cell Painting channels and composite image (all channels overlayed) for each *NF1* genotype. The scale bar represents 25 μM. (**B**) Image analysis and data processing pipeline to derive morphology signature of *NF1* genotype.

Following single-cell QC filtering, we imaged and processed a total of 10,900 *NF1*^*-/-*^ (C04) and 9,780 *NF1*^*+/+*^ (A3) Schwann cells (20,680 total cells) (**Figure 2A**). We collected these cells across three 96-well plates (**Supplementary Figure 3A**; for plate maps, see **Supplemental Figure 4**). We applied a dimensionality reduction method called Uniform Manifold Approximation and Projection (UMAP)^38^ to the single-cell morphology features. We found that single cells were homogeneously clustered across both genotypes, indicating that if an *NF1* genotype signature was present, it manifests in only a few, subtle cell morphology features (**Figure 2B**). We did not observe clustering based on plate, indicating minimal to no batch effects (**Supplementary Figure 3B**). We aggregated the single-cell morphology features to the well-level of the plate, in which each well represents a median profile for all single cells. We calculated Pearson’s correlations by comparing wells with either the same or different *NF1* genotypes. The mean correlation for the same genotype was *r*=0.24, and the different genotype was *r*=0.16. We performed a two-sample t-test, and observed significantly different distributions (t=14.13, p=6.78e-45) (**Figure 2C**). We observed similar differences when looking at individual plates (**Supplementary Figure 3C**). Taken together, these results suggest subtle morphological differences between *NF1* genotypes at both the single-cell and well-population levels. We next demonstrated the ability to detect these differences by training a machine-learning model to classify single-cell *NF1* genotypes.

**Figure 2.**
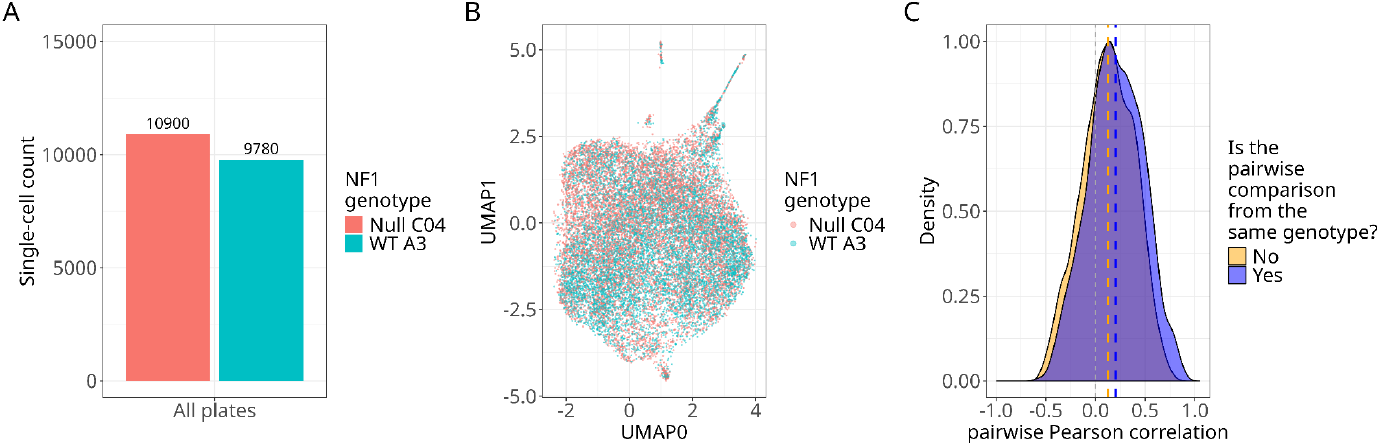
Single-cell and well-level *morphology comparisons between isogenic ipn02*.*3 2λ Schwann cell NF1 genotypes show homogeneity, which suggests subtle morphology differences*. (**A**) We acquired and processed a total of 20,680 single Schwann cells for downstream analysis (e.g., machine learning). (**B**) Uniform Manifold Approximation and Projection (UMAP) applied to the morphological features of all single cells demonstrates homogeneity across genotypes. (**C**) Density plot of Pearson correlations of aggregated single cells at the well level shows similar distributions but a higher mean correlation for the same *NF1* genotype.

### Predicting *NF1* genotype in isogenic Schwann cells with machine learning

We split the dataset into training and testing (data the model has never seen) sets using a manual method that maximizes the number of single cells per genotype in each data split while maintaining an equivalent number of cells per genotype in each set (see methods for details). This procedure was necessary to ensure our model did not overly emphasize learning any specific genotype. We trained a penalized logistic regression classifier to predict *NF1* genotype of single Schwann cells, which involved a procedure known as cross-validation to select an optimal model configuration (see methods for details). To compare the model’s performance, we created a negative control baseline by independently randomizing morphology features (shuffled baseline), which would indicate any artificial biases. We assessed model performance on the training, cross-validation, and testing sets. Precision-recall curves showed strong performance in predicting *NF1* genotype in the testing set and were much stronger than the shuffled baseline (**Figure 3A**). Confusion matrices also showed high performance (specificity=0.79, sensitivity=0.78; **Figure 3B**). Lastly, we observed 79-85% accuracy in predicting *NF1* genotype (**Figure 3C**).

**Figure 3.**
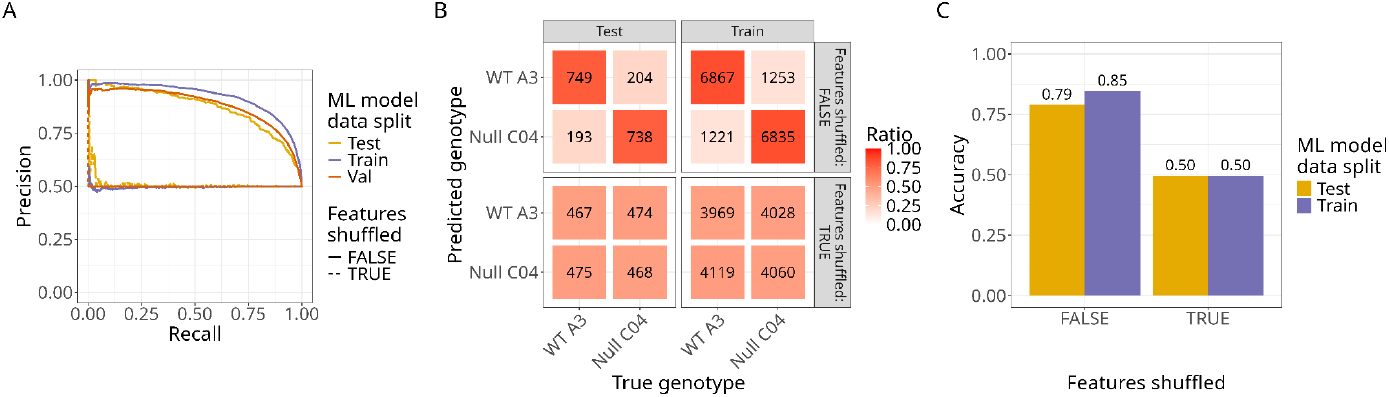
Logistic regression model predicts NF1 genotype in ipn02.3 2λ Schwann cells with high performance. (**A**) Precision-recall curves comparing the final model applied to shuffled (dashed line) and non-shuffled data (solid line). Applying the model to the shuffled data performed worse than the non-shuffled data, demonstrating a strong biological signal between *NF1* genotypes. (**B**) Confusion matrices from the training and testing sets show higher performance across genotypes in non-shuffled data compared to shuffled data. (**C**) Accuracies demonstrate high performance in predicting both genotypes from the training and testing sets compared to shuffled data.

### Multiple organelle morphologies are impacted by *NF1* genotype in Schwann cells

CellProfiler outputs interpretable morphology features, including intensities, textures, granularities, areas, and shapes per organelle.^28^ The machine learning model learned from 894 morphology features and assigned weights, or importance scores, per morphology feature. This combination of morphology features represents a high-dimensional signature of how the *NF1* genotype influences cell morphology in otherwise isogenic Schwann cells (**Figure 4**).

**Figure 4.**
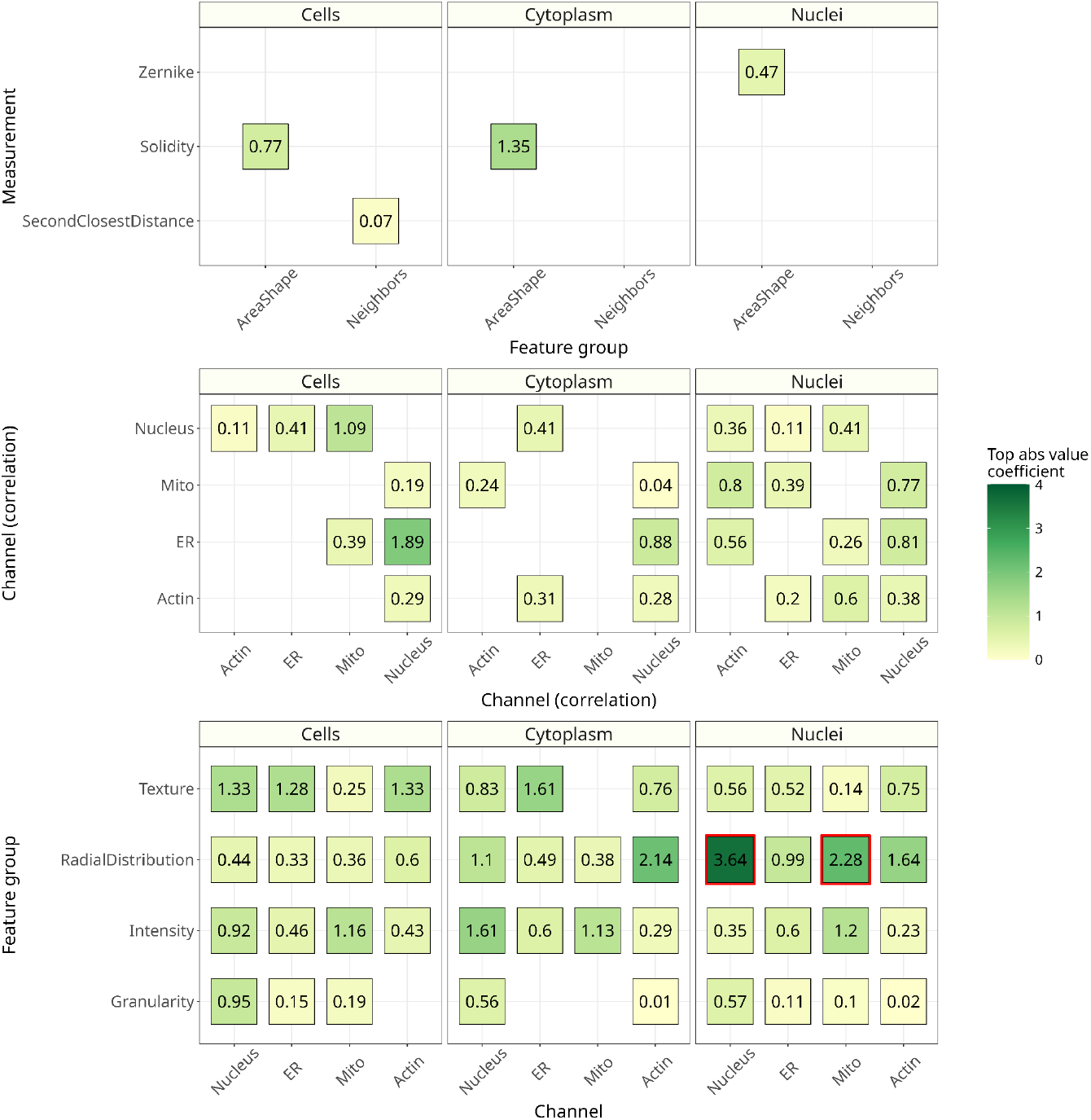
Significant cell morphology features come from multiple kinds of measurement and across organelles. Top absolute value coefficients demonstrate which measurements/organelles/compartments are important in determining the differences between *NF1* genotypes. Not one feature is important in this prediction; many features in our models are important for making accurate genotype predictions. The red boxes indicate the two most important features based on absolute value. Before absolute value, the model coefficients range from positive to negative, where positive indicates more importance in predicting the *NF1* WT A03 (*NF1*^+/+^) genotype and vice versa. Of the two top features, both are highest in the positive range before absolute value. The highest coefficient is the distribution of the nuclei stain on the edge of nuclei, and the second highest coefficient is the distribution of the mitochondria stain on top of nuclei.

Different cell morphology features are important for distinguishing *NF1* genotypes in Schwann cells. The two most important feature categories were the distribution of the nuclei stain on the edge of nuclei (**Figure 5A**) and the distribution of the mitochondria stain on top of nuclei (as they are imaged from above) (**Figure 5B**).

**Figure 5.**
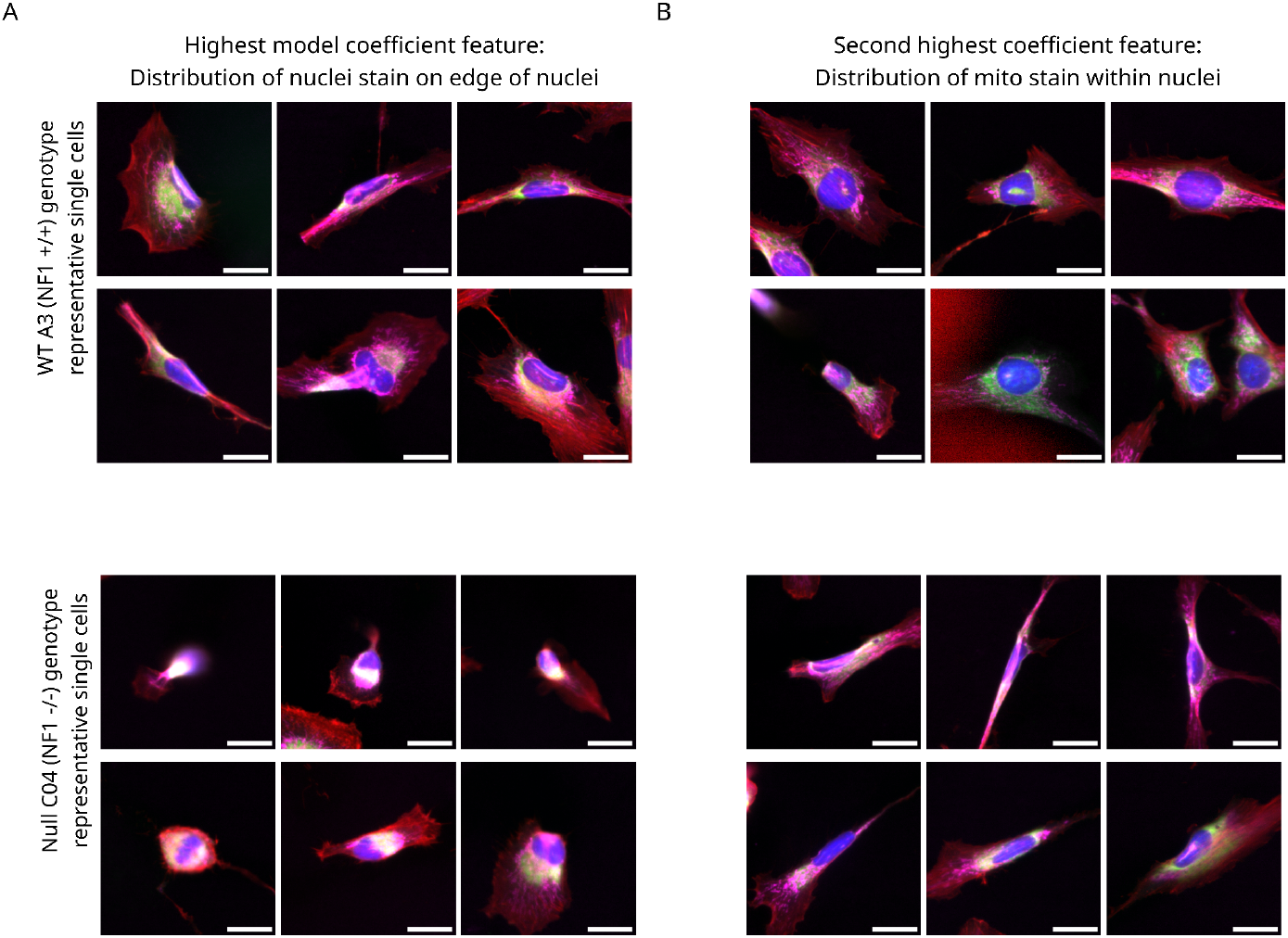
Image montages of the top two most important features in the machine learning model to visualize Schwann cell morphology differences between NF1 genotypes. (**A**) The image montage of representative single cells per *NF1* genotype for the highest coefficient from the machine learning model, which is the distribution of the nuclei staining at the edge of nuclei. As this feature is higher in *NF1* WT A3 (*NF1*^+/+^) cells, we show *NF1* WT A3 single cells with the highest feature values and the lowest feature values for the *NF1* Null C04 (*NF1*^-/-^). (**B**) The image montage of representative single cells per *NF1* genotype for the second highest coefficient from the machine learning model, which is distribution of the mitochondria stain on top of the nuclei. .s this feature is higher in *NF1* WT A3 (*NF1*^+/+^) cells, we show *NF1* WT A3 single cells with the highest feature values and the lowest feature values for the *NF1* Null C04 (*NF1*^-/-^). Both image montages show composite single-cell images with a scale bar of 25 μM. The colors of the images represent each organelle: blue=nucleus, green=endoplasmic reticulum, magenta=mitochondria, and red=F-actin.

### Our machine learning model does not generalize to an additional isogenic pair of Schwann cells

We evaluated the model’s performance on a new “holdout” plate (single cells the model has never seen before) containing an additional isogenic pair (*NF1*^*+/+*^ *and NF1*^*-/-*^ clones) derived from the same ipn02.3 2λ parental cell line. The new plate contained cells from the “original” clones (*NF1* WT “A3” and *NF1* Null “C04”) and these new “derivatives” (e.g., *NF1* WT GFP3, *NF1* Null C23) (**Figure 6A**). We applied the same image analysis pipeline to extract and process 894 single-cell morphology features from all cells. We generated UMAP embeddings and observed that the original and derivative of the ipn02.3 2λ cell line clustered distinctly, indicating stark morphology differences between the two populations (**Figure 6B**). We applied our original pretrained machine learning model to all single cells and generated precision-recall curves for each cell line population. We found high performance in predicting the original clones, but no better than random chance predictions for the new derivatives (**Figure 6C**). We calculated balanced accuracy scores for each clone, and observed low scores for the derivative (**Figure 6D**). Lastly, to evaluate the morphology difference between the two pairs, we performed a Kolmogorov–Smirnov (KS) test for each of the 894 features. This test quantifies on a scale of 0 to 1 how different a feature is between the two populations, with 1 indicating a complete, non-overlapping difference for all cells. We found many feature differences, including some that were important for the model (**Figure 6E**). We speculate these morphology differences are driven by cell line source, single-cell sorting differences prior to cloning, and differences in *NF1* exon CRISPR locus choice (see methods for complete details). We observed morphology differences between clones of the same *NF1* genotype, which suggests strong contributions of non-biological factors (**Supplemental Figure 5**). In the future, we plan on generating a Schwann cell morphology atlas with various Schwann cell lines (including primary lines), *NF1* mutations, and derivation strategies (e.g., immortalization), which will help us to refine our Schwann cell morphology signature of *NF1* activity.

**Figure 6.**
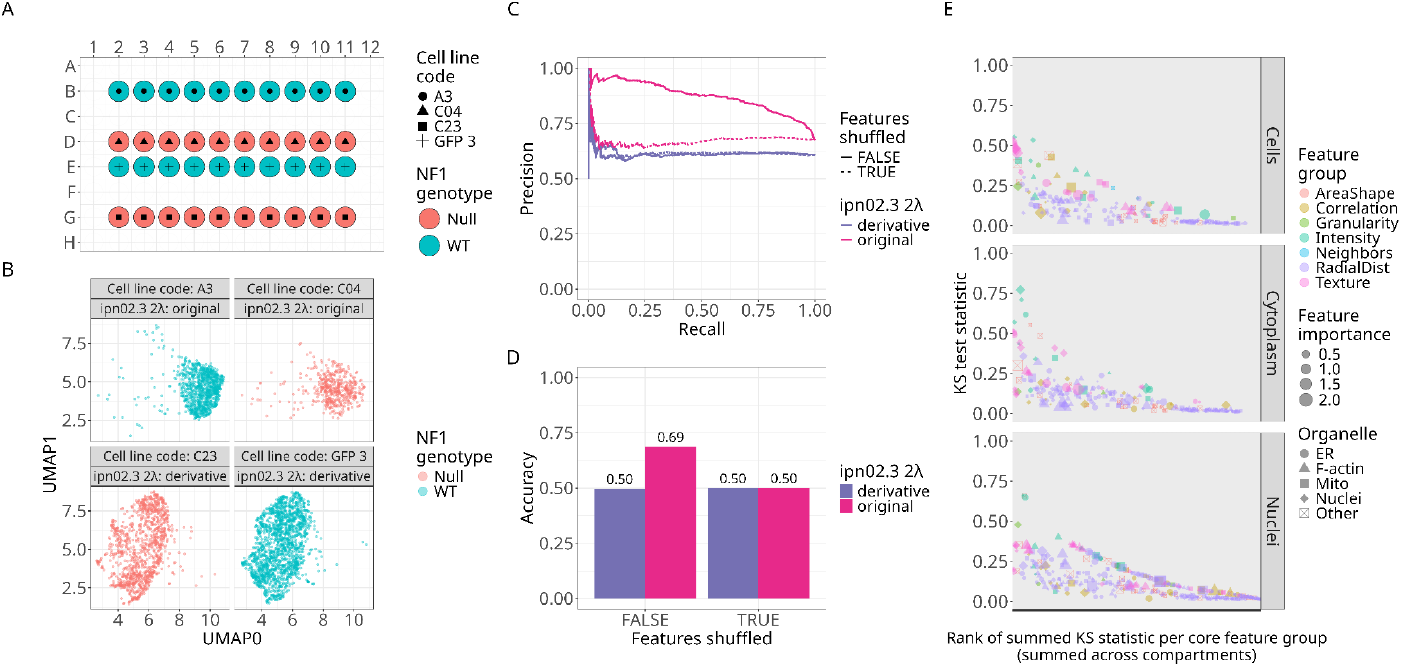
Investigating a new pair of Schwann cells isogenic for NF1. **(A)** Plate map of the new dataset, which includes two newly derived isogenic variants of the ipn02.3λ *NF1* WT (*NF1*^*+/+*^; C23) and Null (*NF1*^*-/-*^; GFP3); see Methods for details. **(B)** UMAP visualization of the 894-feature space comparing the two ipn02.3λ cell line derivatives. The derivative clusters distinctly from the original isogenic pair. **(C)** Precision-recall curves for each isogenic pair show high performance for the original, but no better than random chance predictions for the new derivative. **(D)** Balanced accuracy scores for each isogenic pair. **(E)** Kolmogorov-Smirnov (KS) test statistics applied to each morphology feature comparing the original and new derivative. Many features show significant differences across morphology categories, demonstrating that high-content microscopy can detect subtle variations introduced by different derivation strategies.

## Discussion

Our proof-of-concept experiment demonstrates the feasibility of predicting *NF1* wildtype and null genotypes in Schwann cells using high-content microscopy and machine learning. We achieved balanced accuracy scores of 0.85 for training and 0.79 for testing/holdout data. This performance was substantially better than that of the shuffled datasets, indicating that *NF1* genotype meaningfully influences Schwann cell morphology even after adjusting for increased proliferation in *NF1*^*-/-*^ Schwann cells. These differences are impossible to spot by eye, but we demonstrate that machine learning can detect much more subtle, potentially more important, differences.

Our analysis revealed that a multitude of features from various organelles contributed to the model’s predictive accuracy. This indicates that the phenotypic markers for *NF1* genotypes are described not only by a few features but also by a complex group of multiple cellular characteristics. Specifically, the nucleus, mitochondria, and F-actin cytoskeleton were strongly impacted by *NF1* genotype. For the nucleus, there is a much higher intensity stain on the edge of wildtype Schwann cells. The distribution of mitochondria stain on top of nuclei was also a key predictor of *NF1* genotype. This suggests that the loss of neurofibromin expression leads to differences most strongly in these organelles/subcellular components, but we observed broad and systemic changes across cell structures. In previous experiments, fluorescent staining of neurofibromin has implicated differential localization of F-actin or microtubules^39,40^ and active transport into the nucleus.^41^ Our model corroborates these observations but also quantifies many other morphology differences worthy of follow-up experiments.

Quantitative readouts enhance our ability to collect and interpret high-content microscopy data, which is crucial for identifying potential drug treatments from large-scale drug screens. Previous NF1-focused drug screens focused on targeting *NF1* deficient cells have utilized simpler readouts, such as cell death or viability, by measuring the AC_50_ or IC_50_ or a specifically targeted readout, such as changes in p-ERK expression.^18–20^ However, phenotypic drug screens can measure a thousand-fold more information, including general cell state^42^, drug toxicity^43^, and DNA damage.^42^ This approach can identify compounds that impact cells phenotypically, going beyond identifying compounds that simply kill cells or change single markers. There have been many applications of machine learning using traditional high-content microscopy approaches (which aggregate single cells into bulk profiles), including predicting drug sensitivity^44^ and cell health phenotypes.^42^ Recently, single-cell applications have emerged, demonstrating the potential to predict complex phenotypes, such as predicting MitoCheck phenotypes using single-cell nuclear morphology^37^ and predicting between pyroptotic and apoptotic cell death states.^45^ Based on our findings, we conclude that single-cell high-content microscopy holds strong potential to advance both the study and treatment of NF1. However, before this approach can be applied to phenotypic drug screening, we must further refine the NF1-associated morphology signature in Schwann cells to ensure accuracy and robustness in identifying compounds that restore a healthy, wild-type phenotype in NF1 patient Schwann cells.

Our study has some limitations. First, our assay does not include a heterozygous (*NF1*^+/−^) genotype, limiting our ability to model the full spectrum of neurofibromin production. As our primary goal was to establish proof of concept, we plan to incorporate heterozygous Schwann cells in future studies. Additionally, we derived the cells used to train our machine learning model from a single donor, using an artificial system of hTERT immortalization and isogenic *NF1* knockout via CRISPR-Cas9. Our validation experiments highlighted how critical these technical methods are. We obtained additional wild-type and null cell lines from the same parental cell line, but different technical protocols led to distinct morphological profiles. These findings reinforce that technical variation can introduce significant morphological heterogeneity. While our current study was limited in assessing generalizability in one other cell line, our results indicate that robust machine learning models will require training on a larger and more diverse set of patient-derived cell lines and protocols to minimize the impact of technical variability and improve predictive accuracy.

In the future, we will refine our *NF1* morphology signature in Schwann cells. We plan to build a comprehensive Schwann Cell Morphology Atlas by expanding our dataset to include a range of donors, *NF1* genotypes, derivation protocols, and immortalization strategies. This will enable the development of a high-performing machine learning model that more precisely captures the phenotypic consequences of altered neurofibromin levels. A refined signature will enable us to conduct a phenotypic drug screen using both *NF1*^−/−^ and *NF1*^+/−^ Schwann cells to identify compounds that shift *NF1*-deficient cells toward a healthier morphology. Our methodology can also be extended to other NF1-relevant cell types, including mast cells, fibroblasts, oligodendrocytes, melanocytes, and bone cells. We are also exploring a modified Cell Painting assay that incorporates a fluorescent channel for neurofibromin staining, providing single-cell ground truth of neurofibromin abundance, something not feasible with bulk assays like western blotting. Through this work, we are laying the groundwork for using high-content microscopy to discover novel therapeutic agents with the potential to reverse or slow growth of neurofibromas, offering new treatment strategies for NF1 patients and their families.

## Methods

### Cell Lines

We derived an isogenic *NF1* wildtype (*NF1*^+/+^) and *NF1* null (*NF1*^-/-^) cell line from the hTERT-immortalized human Schwann cell line ipn02.3λ.^1^ We started by isolating single-cell clones of the parental line by sorting with the BD FACSAria III (BD Biosciences). From a monoclonal derived cell line (which we refer to as “A3”), we applied CRISPR-Cas9, targeting the *NF1* gene immediately after the initiator methionine in exon 1.^2^ We confirmed *NF1* mutation and neurofibromin protein loss in our monoclonal knockouts by sequencing and western blots. We call the isogenic wildtype line “A3” and the isogenic *NF1* null line “C04”. For validation experiments (see **Figure 6**), we used another set of clones, previously derived, also from the ipn02.3 2λ Schwann cell line, described in Stevens et al. 2025.^3^ For this set of clones, the isogenic wildtype line is referred to as “GFP3” (since these were subjected to CRISPR/Cas9 transfection with an sgRNA targeting GFP) and the isogenic *NF1* null line as “C23” (with *NF1* mutations in exon 3). The only differences between the two sets of clones (A3/C04 vs. GFP3/C23) is the parental cell line source, sorting, and *NF1* exon targeting: The A3/C04 pair were derived from ipn02.3λ, purchased from American Type Culture Collection (ATCC; CRL-3392), while the GFP3/C23 pair were derived from ipn02.3λ gifted by Dr. Peggy Wallace (University of Florida). While the A3/C04 pair were derived from an ipn02.3λ parental line using a high-pressure, FACS-based sorting, the GFP/C23 did not experience this sorting selection. Lastly, we targeted *NF1* exon 1 in the A3/C04 pair and *NF1* exon 3 in the GFP3/C23 pair. We maintained all cells in DMEM supplemented with Glutamax and 10% FBS.

### Cell painting and image acquisition

We plated cells of each *NF1* genotype in 96-well black, clear bottom plates (Greiner) in either 10% FBS (plate A) or 5% FBS (plates B and C) in DMEM supplemented with Glutamax. We cultured cells for 24 hours at 37°C, 5% CO2. For mitochondria staining, we treated cells with MitoTracker for 30 minutes prior to fixation. After mitochondria staining, we fixed and permeabilized cells, and treated them with the Image-iT Cell Painting kit (Thermofisher). We performed a modified Cell Painting assay, excluding staining for cytoplasmic RNA, nucleoli, and Golgi. Specifically, we used Hoechst (nuclei), 488-tagged Concanavalin A (ER), MitoTracker® Deep Red (mitochondria), and 568 Phalloidin (F-actin). We acquired gray-scale, 16-bit images per FOV using a four-channel plate reader (Biotek, Cytation5). We used 48 wells in plates A and B, with twelve replicate wells per seeding density (500, 1000, 2000, and 4000), with six replicates of each *NF1* isogenic cell line. Plate C utilized 48 wells, with 24 replicate wells per *NF1* isogenic cell line. Plate D utilized 60 wells, with 10 replicate wells per NF1 isogenic cell line.

### Whole image quality control

We assessed the quality of the images from all plates to remove those of poor quality (**Supplemental Figure 6**). Poor-quality images included large smudges and/or debris, out-of-focus organelles, and over-saturated organelles. We utilized CellProfiler’s image quality metrics to decide two image quality features best distinguished between poor and good quality images: Power Log Log Slope (to represent blur) and Percent Maximal (to represent saturation). These features were recommended by Caciedo et al. in their paper describing strategies for image-based profiling.^33^ We observed that, considering all images, the blur metric forms a Gaussian distribution, where poor-quality images are found at the distribution tails. We set the threshold for out-of-focus images in our CellProfiler pipeline as two standard deviations above and below the mean (this process categorized 95% of the data as good quality). As for saturation, we observed a distribution skewed towards 0 but with a long tail. We set the threshold as two standard deviations above the mean, as any values above are likely oversaturated and of poor quality. After quality control, 968 image sets failed QC (4.7%), and we did not process cells from these images. Note that our procedure for whole image quality control is different from our procedure for single-cell quality control, which occurs after image analysis.

### Image analysis

We used CellProfiler v4.2.4, a standard software in the image analysis field, to perform our image analysis pipelines.^47^ We applied an illumination correction pipeline for each image set (a group of channels per FOV) that passed quality control. This pipeline created an illumination correction function for each channel per image set, applied the function, and saved each corrected image set as TIFF files. Illumination correction smoothens the pixel intensity of the image, ensuring that our single-cell analysis is not biased based on the cell’s location in the image.

We next applied a segmentation and feature extraction pipeline using CellProfiler. We segmented three compartments for each single cell per image set: nuclei, cells, and cytoplasm. We derived nuclei segmentation using the DAPI channel, cell segmentation from the RFP channel (seeding the nuclei segmentation as the input to identify cells), and cytoplasm segmentation by subtracting the nuclei from the cell segmentations. These three segmentations are represented by binary masks (0/1) that we used to extract features from each compartment for every single cell while ignoring the images’ background. We extracted a total of 2,313 features across all compartments per single cell. The pipeline outputs an SQLite database per plate, with each compartment as a table, with all features extracted.

### Image-based profiling

To perform downstream analysis and machine learning, we processed the CellProfiler output with the standard image-based profiling pipeline. First, using CytoTable^34^, we converted the SQLite databases into parquet files. We merged each compartment table based on object number, which leaves one single cell per row and all morphology features as columns. We next used pycytominer,^35^ a bioinformatics software tool for processing image-based profiles, to complete the processing. First, we performed annotation where we combined metadata per well from a plate map with every single cell (e.g., genotype, plate, etc.). After annotation, we normalized the cell morphology features using the standard scalar method, which transforms data to zero mean with unit variance. We calculated standard scalar normalization based on the whole plate for Plates A, B, and C. For Plate D, we fit the standard scaler using only the Null (C04) and WT (A3) cell lines (and subsequently normalized the full plate). This procedure allows us to directly compare Plate D with the original three plates. Lastly, we performed feature selection, which removes non-informative and redundant features. We used four operations in feature selection: variance threshold, correlation threshold, blocklist, and drop NaN columns. We set the variance threshold as the default frequency cutoff of 0.05, which will remove features with low variance across all single cells. We set the correlation threshold as the default Pearson correlation threshold of 0.9, where we removed one of the highly correlated features by selecting the feature with the lowest absolute value correlation to all the other features. We removed features included in the blocklist default file as set by pycytominer. Lastly, we removed features with at least one NaN value per single cell by setting the na_cutoff parameter as 0. We took the intersection of the features selected across plates as the final feature space for training the machine learning model. This procedure resulted in 907 features in common. After feature selection, we performed aggregation, calculating the median values of each morphology feature for every single cell per well. We segmented 11,286 single cells from 48 wells in Plate A, 5,506 single cells from 48 wells in Plate B, 5,793 single cells from 48 wells for Plate C, and 7,383 single cells from Plate D. We use the well-level post-feature selection profiles to assess correlations between genotypes. We used single-cell profiles for machine learning. Analyzing the single-cell profiles include all heterogeneity within each well but risk including low quality segmentations, while the aggregated profiles reduces heterogeneity but removes outliers which may skew single-cell interpretation.

### Quality control of single-cell profiles

A critical step in our image analysis pipeline is segmenting single cells. Often, this procedure can segment cells incorrectly. Errors include segmenting two overlapping cells as one, segmenting debris rather than single cells, and over/under segmenting a single cell as a result of increased/reduced pixel contrast. To reduce the noise of including poor segmentations in our model, we used a single-cell quality control (QC) software called coSMicQC. By utilizing CellProfiler features, we can detect outliers that are related to poor quality segmentations. We used a combination of two features; solidity, which measures irregularity in shape, and total intensity. We removed cells with 1.25 standard deviations below the mean for nuclei solidity and 2 standard deviations above the mean for the intensity, using both features combined. This combination removes poorly segmented cells where two nuclei near each other are segmented as one (**Supplemental Figure 2; top**). Additionally, because we expect *NF1*-null Schwann cells to exhibit increased proliferation^8^, which could bias the machine learning model to associate the *NF1*^-/-^ genotype with a higher prevalence of mitotic cells, we used coSMicQC to identify and remove mitotic cells prior to analysis. Based on our prior work^7^, we used two intensity features, upper quartile intensity (how bright the pixel intensity is within nuclei) and MAD intensity (how variable the pixel intensity values are within nuclei) (**Supplemental Figure 2; bottom**). We removed cells with 2 standard deviations above the mean of each feature independently. After detecting outliers, coSMicQC removed approximately 9.2% of single-cells across all plates.

### Machine learning: Data splitting and sampling

To avoid bias toward our model learning either *NF1* genotype preferentially, we balanced the distribution of genotypes in our training and testing sets. Additionally, we prevented the training and test set to include cells from the same well. By preventing the influence of neighboring cells leaking from the training set into the test set, we are able to effectively evaluate the generalizability of our machine learning model to new cell populations. We manually split training and test wells to achieve an approximate ratio of 90% training cells to 10% testing cells. We then randomly downsampled the training and test sets to achieve an equal number of cells per genotype. This procedure avoids the pitfalls of evaluating a supervised machine learning model trained on data with class imbalance.^48^

### Machine learning: Model training and evaluation

We developed an interpretable machine-learning approach to identify important morphology features responsible for discerning *NF1* genotypes. We trained a machine learning model to predict NF1 genotype from single-cell morphology features using L2-regularized logistic regression. L2 regularization reduces overfitting without encouraging sparsity of morphology weights after feature selection.^49^ We determined the optimal inverse regularization strength (λ = *C*^-1^) by performing a random search on the class-balanced training data with a stratified 8-fold cross-validation. We evaluated the accuracy on the holdout cross-validated data in 500 logistic regression models, each with an inverse regularization strength λ ∼ *U*(0, 200), and we chose hyperparameters that maximized cross-validation accuracy. We then computed validation metrics from all cross-validation folds, which have previously been shown to reduce bias.^49^ To compare our results to a negative control baseline, we randomized each feature independently for each dataset, performed the same cross-validation procedure, and then evaluated each of these augmented datasets. After this randomization, we retrained the model on the entire training split using the optimal hyperparameter, and evaluated performance on the training and testing splits.

## Data availability

Datasets containing the original images can be found on figshare at https://figshare.com/projects/NF1_Schwann_Cell_Genotype_Cell_Painting_Assay/161620.^50–54^ Code to download datasets, perform image analysis with CellProfiler, and image-based profiling with CytoTable and pycytominer is available on GitHub at https://github.com/WayScience/nf1_cellpainting_data.^55^ Code to perform all analyses (EDA, machine learning model training, validation, and figure making) is available at https://github.com/WayScience/nf1_schwann_cell_morphology_signature.^56^

## Acknowledgements

We would like to thank Dave Bunten, Mike Lippincott, Erik Serrano, and Roshan Kern for their contribution to code reviews. Thanks to Dustin Rubinstein and Brent Lehman (University of Wisconsin-Madison Biotechnology Center) for engineering of iNFixion cell lines. We also thank Peggy Wallace for donating the ipn02.3λ parental cell line we used for validating isogenic clones.

## Funding

This work was supported by the Department of Defense (DoD) Congressionally Directed Medical Research Programs (CDMRP) Neurofibromatosis Research Program (NFRP) under an Investigator-Initiated Award (HT9425-23-1-0490) to J.A.W. and M.M-H. and an Early Investigator Research Award (EIRA) (HT9425-23-1-0831) to S.J.B. S.J.B. was also funded in part by the National Human Genome Research Institute (NHGRI) under award #T32HG010464.

## Conflicts of interest

MMH and HS are employees of Infixion Bioscience. GPW serves on the scientific advisory board of Infixion Bioscience.

## Notes

https://figshare.com/projects/NF1_Schwann_Cell_Genotype_Cell_Painting_Assay/161620

https://github.com/WayScience/nf1_cellpainting_data

https://github.com/WayScience/nf1_schwann_cell_morphology_signature

### Summary of Updates

We added a new dataset derived from a different clone of the same parental Schwann cell line. Our model did not generalize well to this new cell line, and we present evidence suggesting this is due to differences in the derivation process. Additionally, we implemented a rigorous single-cell quality control pipeline and retrained all machine learning models.

## References

1. Wang, W. et al. Impacts of NF1 Gene Mutations and Genetic Modifiers in Neurofibromatosis Type 1. Front. Neurol. 12, 704639 (2021).

2. Pemov, A., Park, C., Reilly, K. M. & Stewart, D. R. Evidence of perturbations of cell cycle and DNA repair pathways as a consequence of human and murine NF1-haploinsufficiency. BMC Genomics 11, 194 (2010).

3. Johnson, K. J. et al. Development of an international internet-based neurofibromatosis Type 1 patient registry. Contemp. Clin. Trials 34, 305–311 (2013).

4. Legius, E. & Brems, H. Genetic basis of neurofibromatosis type 1 and related conditions, including mosaicism. Childs. Nerv. Syst. 36, 2285–2295 (2020).

5. Rossi, S. et al. Neurofibromin C terminus-specific antibody (clone NFC) is a valuable tool for the identification of NF1-inactivated GISTs. Mod. Pathol. 31, 160–168 (2018).

6. Serra, E. et al. Confirmation of a double-hit model for the NF1 gene in benign neurofibromas. Am. J. Hum. Genet. 61, 512–519 (1997).

7. Colman, S. D., Williams, C. A. & Wallace, M. R. Benign neurofibromas in type 1 neurofibromatosis (NF1) show somatic deletions of the NF1 gene. Nat. Genet. 11, 90–92 (1995).

8. Park, S.-J. et al. Serum biomarkers for neurofibromatosis type 1 and early detection of malignant peripheral nerve-sheath tumors. BMC Med. 11, 109 (2013).

9. Cannon, A. et al. Cutaneous neurofibromas in Neurofibromatosis type I: a quantitative natural history study. Orphanet J. Rare Dis. 13, 31 (2018).

10. Needle, M. N. et al. Prognostic signs in the surgical management of plexiform neurofibroma: the Children’s Hospital of Philadelphia experience, 1974-1994. J. Pediatr. 131, 678–682 (1997).

11. Higham, C. S. et al. The characteristics of 76 atypical neurofibromas as precursors to neurofibromatosis 1 associated malignant peripheral nerve sheath tumors. Neuro. Oncol. 20, 818–825 (2018).

12. Lim, Z., Gu, T. Y., Tai, B. C. & Puhaindran, M. E. Survival outcomes of malignant peripheral nerve sheath tumors (MPNSTs) with and without neurofibromatosis type I (NF1): a meta-analysis. World J. Surg. Oncol. 22, 14 (2024).

13. Gross, A. M. et al. Selumetinib in Children with Inoperable Plexiform Neurofibromas. N. Engl. J. Med. 382, 1430–1442 (2020).

14. Weiss, B. D. et al. NF106: A Neurofibromatosis Clinical Trials Consortium Phase II Trial of the MEK Inhibitor Mirdametinib (PD-0325901) in Adolescents and Adults With NF1-Related Plexiform Neurofibromas. J Clin Oncol 39, 797–806 (2021).

15. Sharawat, I. K., Panda, P. K., Sihag, R. K., Panda, P. & Dawman, L. Efficacy and safety profile of selumetinib in symptomatic inoperable plexiform neurofibromas. J. Neurosurg. Sci. 66, 501–510 (2022).

16. Gross, A. M. et al. Selumetinib in children with neurofibromatosis type 1 and asymptomatic inoperable plexiform neurofibroma at risk for developing tumor-related morbidity. Neuro. Oncol. 24, 1978–1988 (2022).

17. Frost, M. et al. Rationale for haploinsufficiency correction therapy in neurofibromatosis type 1. J. Transl. Genet. Genom. 6, 403–428 (2022).

18. Bouley, S. J. et al. Chemical genetic screens reveal defective lysosomal trafficking as synthetic lethal with NF1 loss. J. Cell Sci. 137, (2024).

19. See, W. L., Tan, I.-L., Mukherjee, J., Nicolaides, T. & Pieper, R. O. Sensitivity of glioblastomas to clinically available MEK inhibitors is defined by neurofibromin 1 deficiency. Cancer Res. 72, 3350–3359 (2012).

20. Maertens, O. et al. Elucidating distinct roles for NF1 in melanomagenesis. Cancer Discov. 3, 338–349 (2013).

21. Vincent, F. et al. Phenotypic drug discovery: recent successes, lessons learned and new directions. Nat. Rev. Drug Discov. 21, 899–914 (2022).

22. Way, G. P., Sailem, H., Shave, S., Kasprowicz, R. & Carragher, N. O. Evolution and impact of high content imaging. SLAS Discov 28, 292–305 (2023).

23. Miller, S. J. et al. Large-scale molecular comparison of human schwann cells to malignant peripheral nerve sheath tumor cell lines and tissues. Cancer Res. 66, 2584–2591 (2006).

24. Ferrer, M. et al. Pharmacological and genomic profiling of neurofibromatosis type 1 plexiform neurofibroma-derived schwann cells. Sci Data 5, 180106 (2018).

25. Suppiah, S. et al. Multiplatform molecular profiling uncovers two subgroups of malignant peripheral nerve sheath tumors with distinct therapeutic vulnerabilities. Nat. Commun. 14, 2696 (2023).

26. Kim, H. A., Rosenbaum, T., Marchionni, M. A., Ratner, N. & DeClue, J. E. Schwann cells from neurofibromin deficient mice exhibit activation of p21ras, inhibition of cell proliferation and morphological changes. Oncogene 11, 325–335 (1995).

27. Bray, M.-A. et al. Cell Painting, a high-content image-based assay for morphological profiling using multiplexed fluorescent dyes. Nat. Protoc. 11, 1757–1774 (2016).

28. Carpenter, A. E. et al. CellProfiler: image analysis software for identifying and quantifying cell phenotypes. Genome Biol. 7, R100 (2006).

29. Caicedo, J. C. et al. Cell Painting predicts impact of lung cancer variants. Mol. Biol. Cell 33, ar49 (2022).

30. Li, H., Chang, L.-J., Neubauer, D. R., Muir, D. F. & Wallace, M. R. Immortalization of human normal and NF1 neurofibroma Schwann cells. Lab. Invest. 96, 1105–1115 (2016).

31. Stevens, M. et al. Inhibition of autophagy as a novel treatment for neurofibromatosis type 1 tumors. Mol Oncol 19, 825–851 (2025).

32. Singh, S., Bray, M.-A., Jones, T. R. & Carpenter, A. E. Pipeline for illumination correction of images for high-throughput microscopy. J. Microsc. 256, 231–236 (2014).

33. Caicedo, J. C. et al. Data-analysis strategies for image-based cell profiling. Nat. Methods 14, 849–863 (2017).

34. Bunten, D., Alquaddoomi, F., Serrano, E., & Way, G. CytoTable. (Github).

35. Serrano, E. et al. Reproducible image-based profiling with Pycytominer. Nature Methods (2025) doi:10.1038/s41592-025-02611-8.

36. coSMicQC: Single Cell Morphology Quality Control (coSMicQC). (Github).

37. Tomkinson, J., Kern, R., Mattson, C. & Way, G. P. Toward generalizable phenotype prediction from single-cell morphology representations. BMC Methods 1, 2024.03.13.584858 (2024).

38. McInnes, L., Healy, J. & Melville, J. UMAP: Uniform Manifold Approximation and Projection for Dimension Reduction. arXiv [stat.ML] (2018).

39. Li, C., Cheng, Y., Gutmann, D. A. & Mangoura, D. Differential localization of the neurofibromatosis 1 (NF1) gene product, neurofibromin, with the F-actin or microtubule cytoskeleton during differentiation of telencephalic neurons. Brain Res. Dev. Brain Res. 130, 231–248 (2001).

40. Gregory, P. E. et al. Neurofibromatosis type 1 gene product (neurofibromin) associates with microtubules. Somat. Cell Mol. Genet. 19, 265–274 (1993).

41. Vandenbroucke, I., Van Oostveldt, P., Coene, E., De Paepe, A. & Messiaen, L. Neurofibromin is actively transported to the nucleus. FEBS Lett. 560, 98–102 (2004).

42. Way, G. P. et al. Predicting cell health phenotypes using image-based morphology profiling. Mol. Biol. Cell 32, 995–1005 (2021).

43. Nyffeler, J. et al. Bioactivity screening of environmental chemicals using imaging-based high-throughput phenotypic profiling. Toxicol. Appl. Pharmacol. 389, 114876 (2020).

44. Kelley, M. E. et al. High-content microscopy reveals a morphological signature of bortezomib resistance. bioRxiv (2023) doi:10.1101/2023.05.02.539137.

45. Lippincott, M. J. et al. A morphology and secretome map of pyroptosis. bioRxiv 2024.04.26.591386 (2024) doi:10.1101/2024.04.26.591386.

46. Zhang, T., Han, T., Dong, Z., Li, C. & Lu, W. Characterization of Two Loss-of-Function NF1 Variants in Chinese Patients and Potential Molecular Interpretations of Phenotypes. Front. Genet. 12, 660592 (2021).

47. Stirling, D. R. et al. CellProfiler 4: improvements in speed, utility and usability. BMC Bioinformatics 22, 433 (2021).

48. Lee, W. & Seo, K. Downsampling for Binary Classification with a Highly Imbalanced Dataset Using Active Learning. Big Data Research 28, 100314 (2022).

49. Forman, G. & Scholz, M. Apples-to-apples in cross-validation studies: pitfalls in classifier performance measurement. SIGKDD Explor. Newsl. 12, 49–57 (2010).

50. Tomkinson, J., Mattson-Hoss, M., Sarnoff, H. & Way, G. Plate 3 and Plate 3’ (prime). 10.6084/m9.figshare.22592890.v1 (2023).

51. Tomkinson, J., Mattson-Hoss, M., Sarnoff, H. & Way, G. Plate 5. 10.6084/m9.figshare.26759914.v1 (2024).

52. Tomkinson, J., Mattson-Hoss, M., Sarnoff, H. & Way, G. Plate 1. 10.6084/m9.figshare.22233292.v1 (2023).

53. Tomkinson, J., Mattson-Hoss, M., Sarnoff, H. & Way, G. Plate 2. 10.6084/m9.figshare.22233700.v1 (2023).

54. Tomkinson, J., Mattson-Hoss, M., Sarnoff, H. & Way, G. Plate 4. 10.6084/m9.figshare.23671056.v1 (2023).

55. Tomkinson, J., Mattson, C. Way, G., Serrano, E. nf1_schwann_cell_painting_data. doi:10.5281/zenodo.13345304.

56. Mattson, C., Tomkinson, J., Way, G. WayScience/nf1_schwann_cell_morphology_signature: v1.1 - Manuscript Updates. doi:10.5281/zenodo.13694800.

