## Supplementary Figures 1-6 for "High-content microscopy and machine learning characterize a cell morphology signature of *NF1* genotype in Schwann cells"

[1]Department of Biomedical Informatics, University of Colorado School of Medicine
[2]Infixion Bioscience
[3]Center for Genomic Medicine, Massachusetts General Hospital

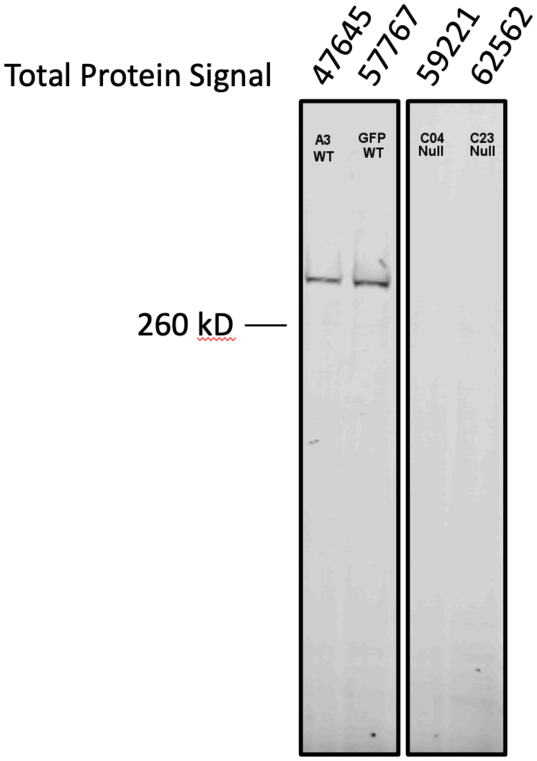

Supplemental Figure 1: Western blots of each cell line demonstrate expected neurofibromin content.

We observe positive neurofibromin signal in *NF1* wild-type (WT) isogenic clones (A3 and GFP3) and the absence of neurofibromin in NF1 Null isogenic clones (C04 and C23). Total protein stain is used to ensure consistent loading.

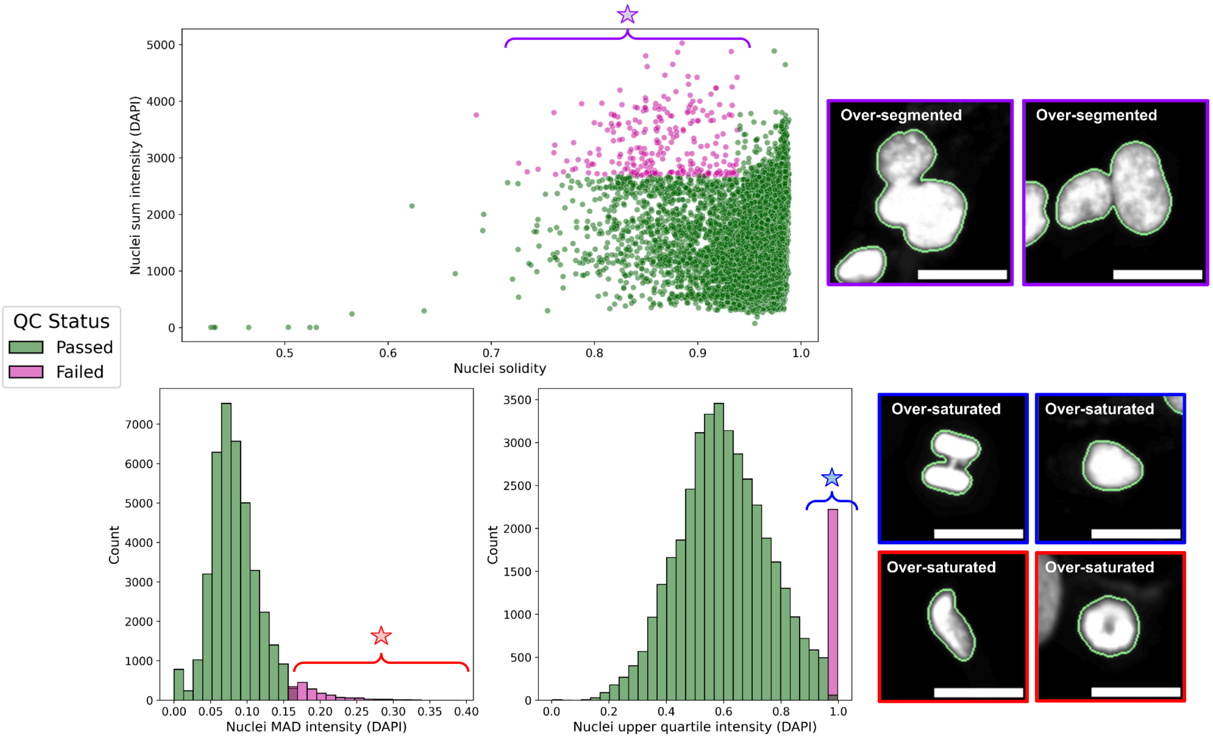

Supplemental Figure 2: Single cell quality control to remote poor quality segmentations and cells undergoing mitosis.

(**top**) We used two CellProfiler features to identify and remove poor-quality segmentations due to technical errors. High intensity and irregular shape features were used together to identify over-segmented nuclei. (**bottom**) We also used two different high-intensity features to identify blurry nuclei and cells undergoing mitosis. The impact of poor-quality nuclei segmentation is minimal within two of the plots, but there is a higher impact in one of the high-intensity (upper quartile intensity) features, based on the histogram plots per plate per feature. Scale bars are 25 μM.

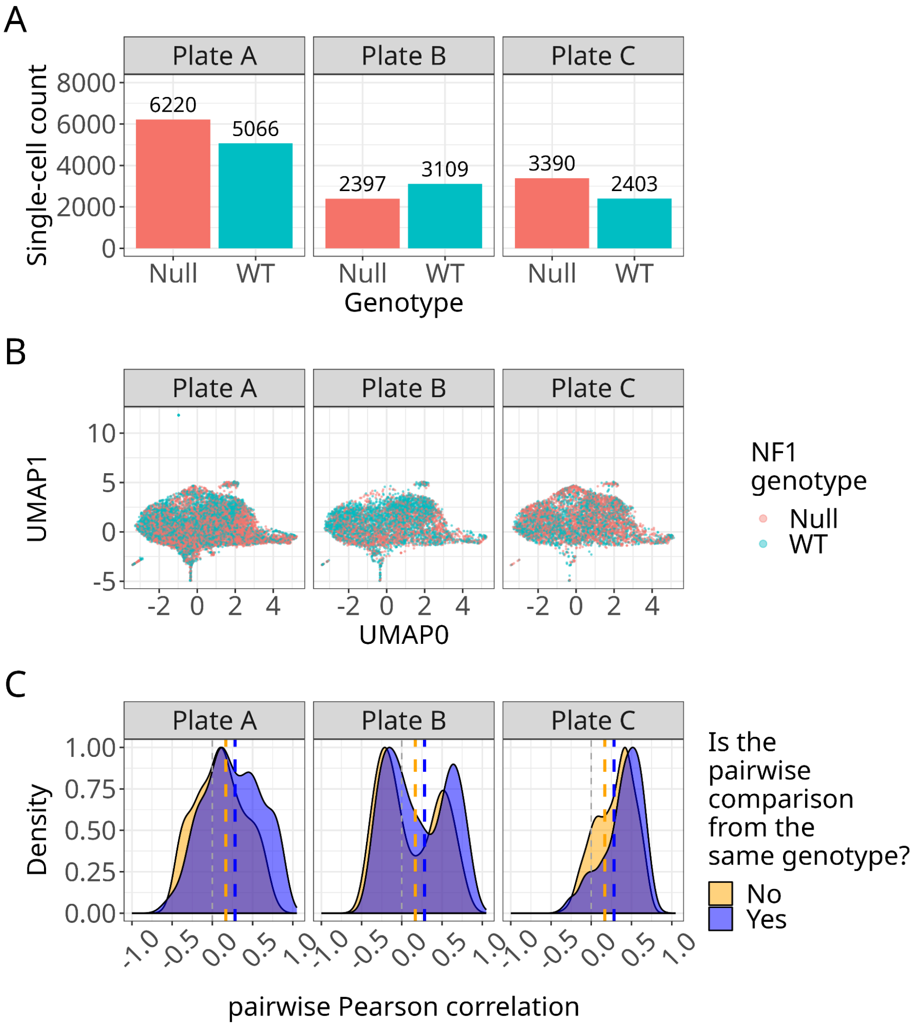

Supplemental Figure 3: Plate-level comparisons of single-cell and well-level morphology

(**A**) Count of single Schwann cells per *NF1* genotype across plates. The cell line is ipn02.3 2λ; the WT cells are called A3, while the null cells are called C04. (**B**) Uniform Manifold Approximation and Projection (UMAP) applied to the morphological feature space split by plate shows no clustering, suggesting no batch effect is present. (**C**) Density plots of Pearson correlations of aggregated single cells at the well level show similar distributions but higher mean correlations for wells of the same genotype.

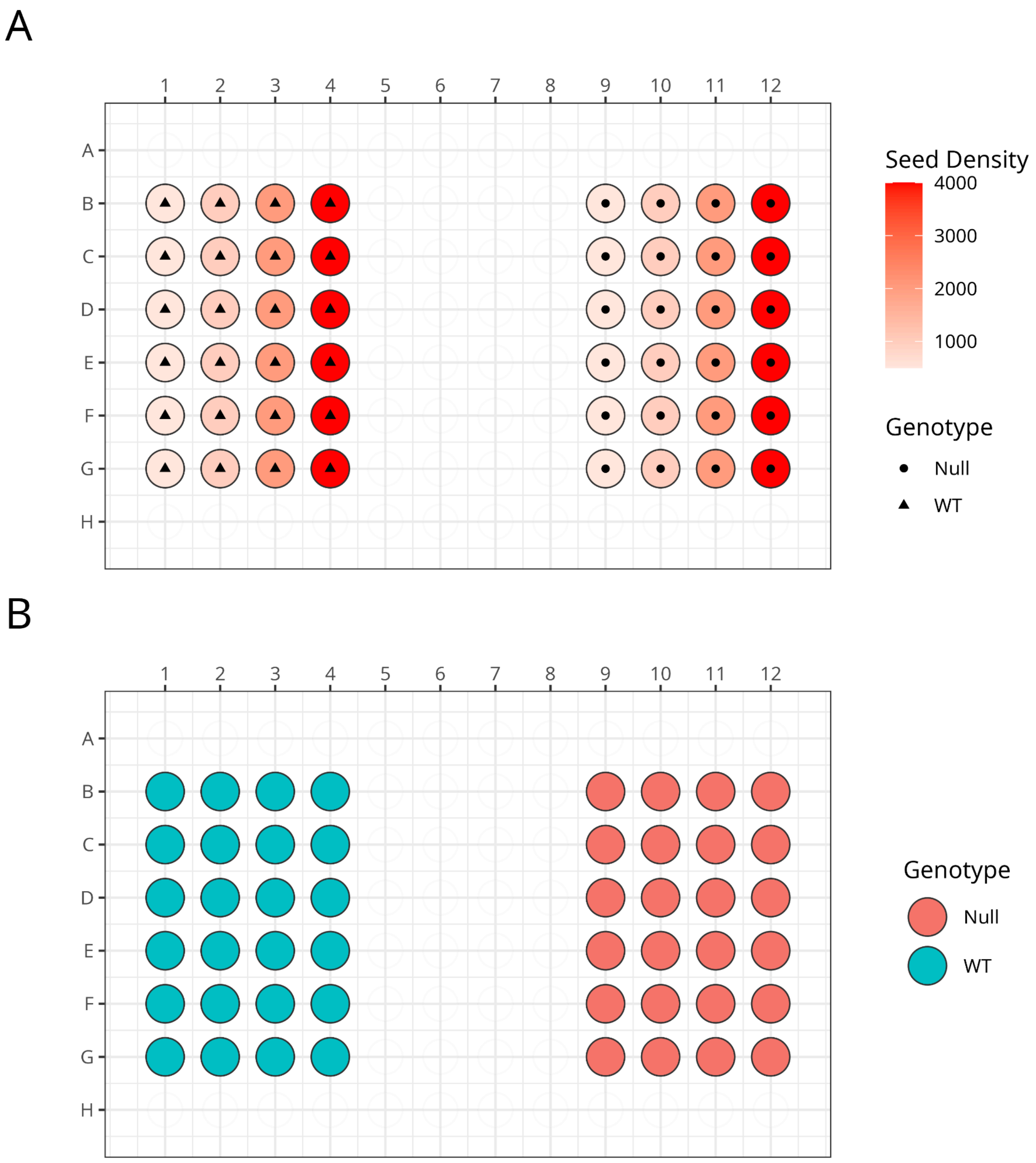

Supplemental Figure 4: Plate map layouts

(**A**) Plate map layout for Plates A and B, with varying seeding densities across two *NF1* genotypes. (**B**) Plate map layout for Plate C with each *NF1* genotype at 1,000 seeding density.

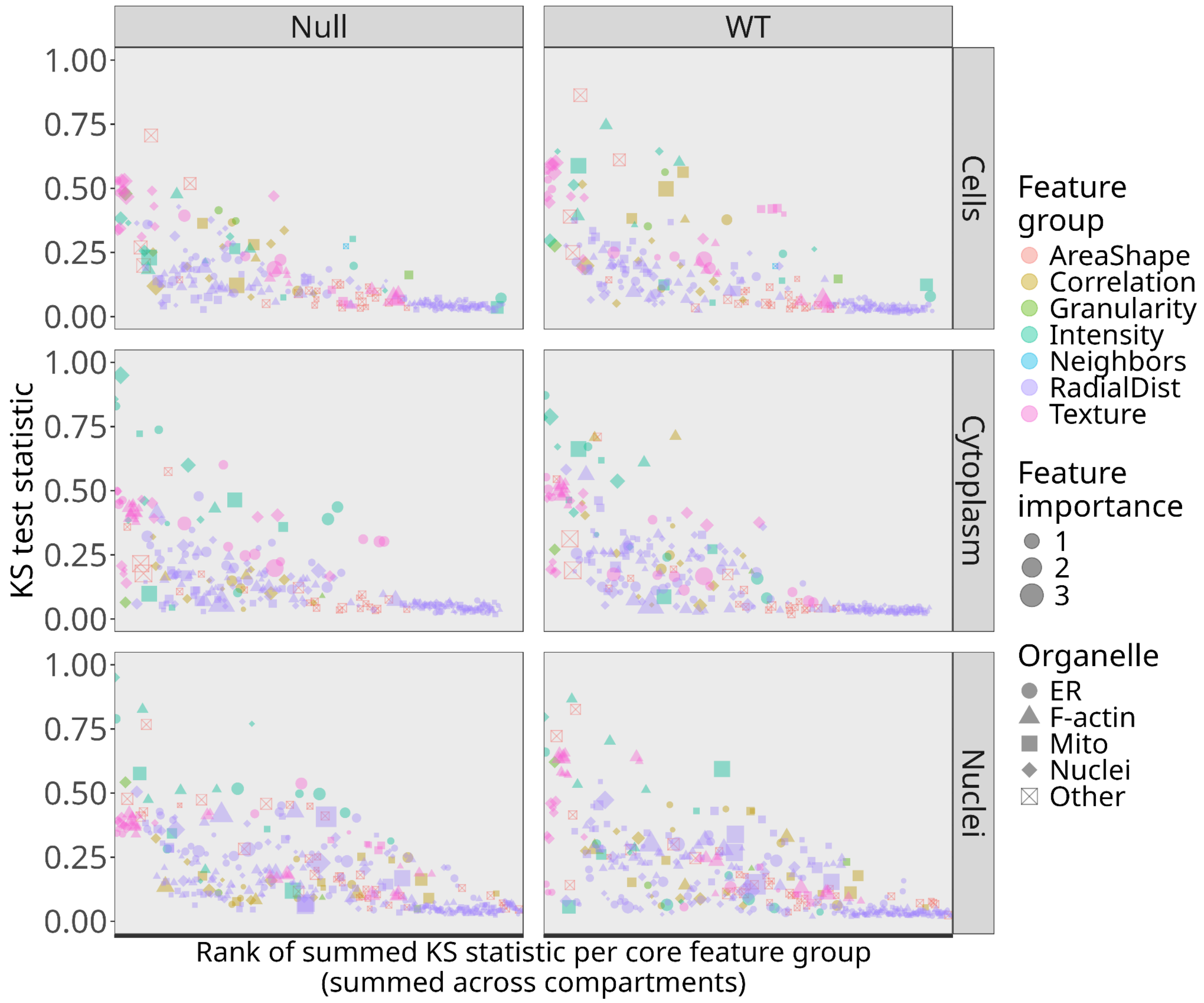

Supplemental Figure 5: KS-test results comparing CellProfiler feature differences between cell lines of the same *NF1* genotype

We performed a Kolmogorov-Smirnov (KS) test comparing across cell line derivatives but within *NF1* genotype. Many features, across measurements and organelles, differ for each genotype. These differences are contributing to the model's lower performance.

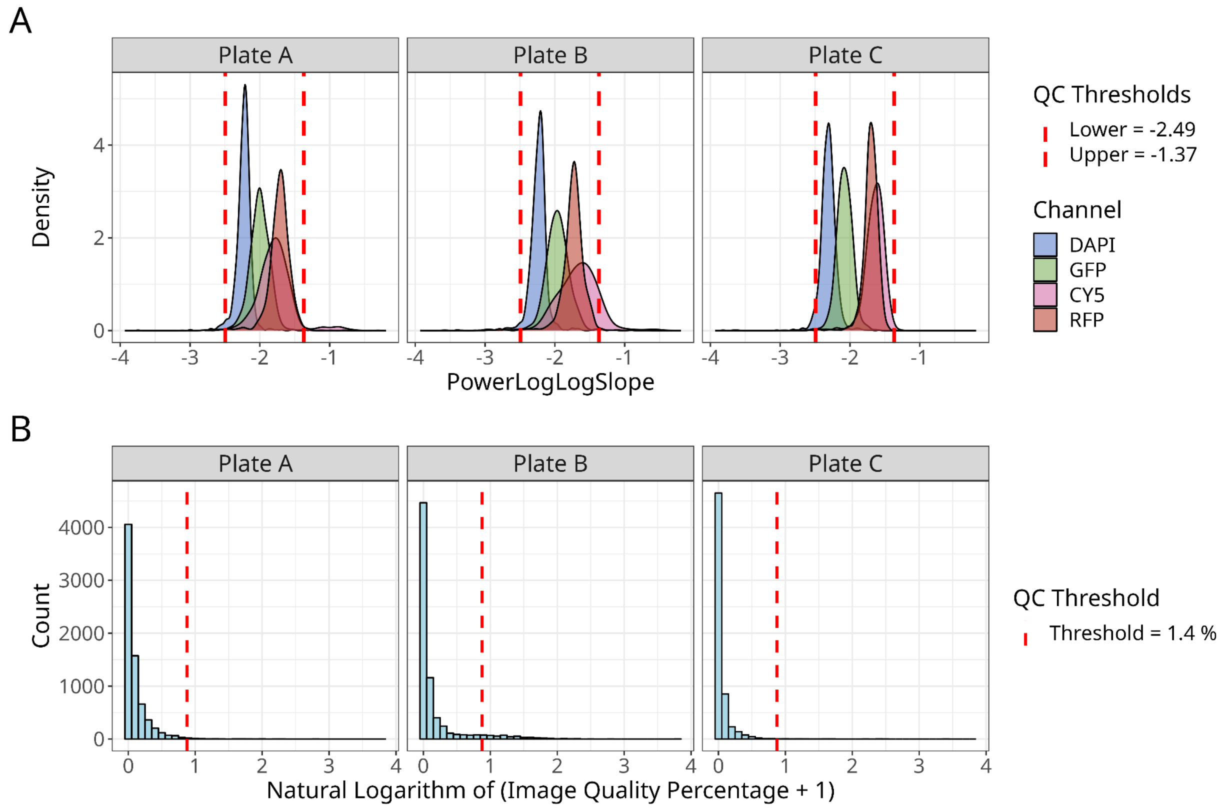

Supplemental Figure 6: Image quality control metrics remove poor-quality fields of view

(**A**) Distribution of PowerLogLogSlope metric, representing blur, per channel across plates shows normal distributions where the tails represent poor quality images. Thresholds (red dotted lines) represent the upper and lower limits of good-quality images that are applied to CellProfiler. (**B**) The distribution of Percent Maximal, representing over-saturation, for all chances across plates shows a right-skewed distribution where most good quality images have a low percent of maximum pixel values. The values have been updated with the natural logarithm plus one for better visualization. The red dotted line represents the threshold at which the images with over 1.4% of pixels at maximum value are considered poor quality.
